## Supplementary Figures for "Leading Basic Modes of Spontaneous Activity Drive Individual Functional Connectivity Organization in the Resting Human Brain"

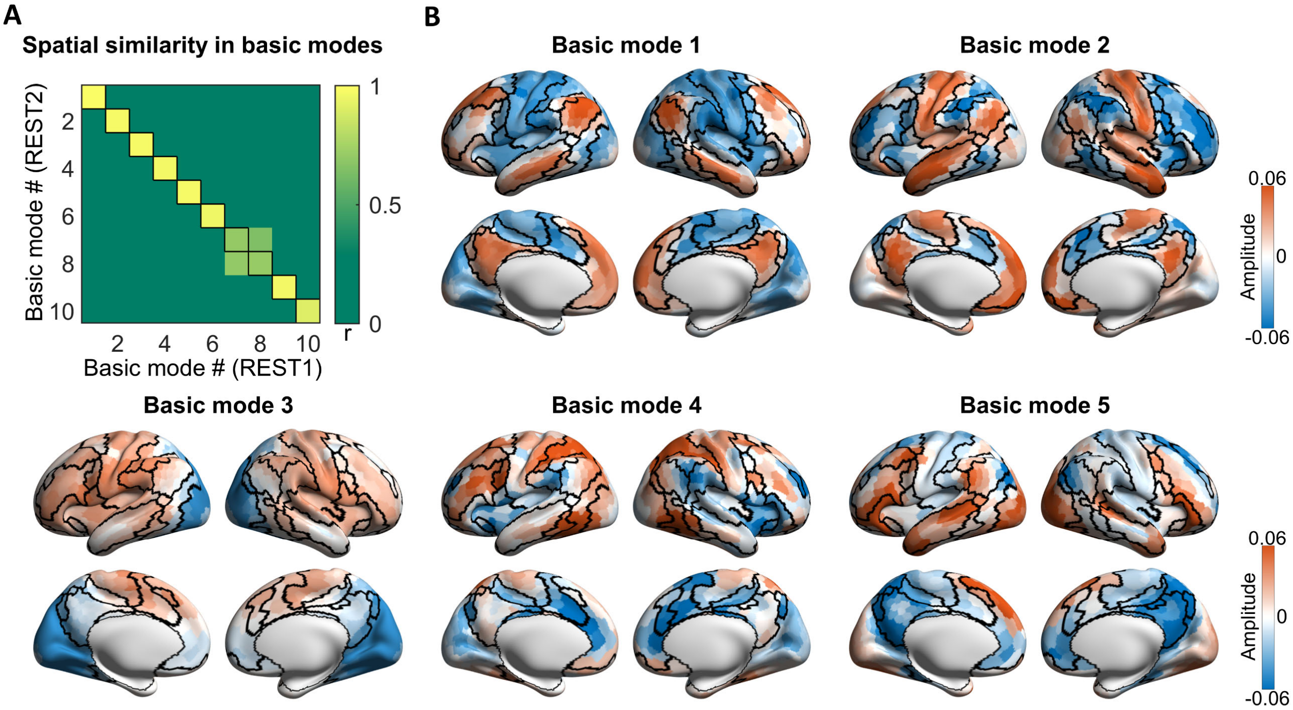


**Figure S1.** Reproducibility of five leading basic modes in REST2 of the HCP dataset. (A) Spatial similarity of the first ten basic modes between two runs (i.e., REST1 and REST2). We identified the basic modes from the REST1 and REST2 of the HCP dataset separately. The first ten basic modes showed an exact correspondence, except for the inversion of the 7th and 8th basic modes. (B) Spatial patterns of the first five basic modes (i.e., leading basic modes) for REST2. Black curves denote the boundaries of the prior seven functional systems[1]. HCP, Human Connectome Project.

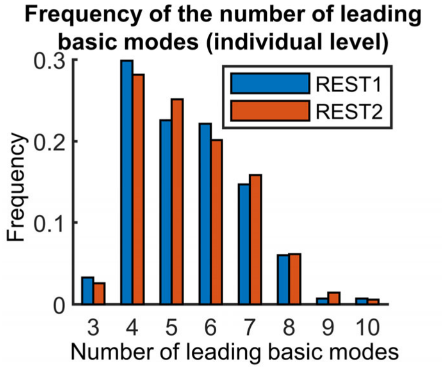


**Figure S2.** Normalized histograms of the number of leading basic modes at the individual level (REST1 and REST2 of the HCP dataset). For each participant, the number of leading basic modes was determined by identifying the elbow point in the weight curve using the Kneedle algorithm[2]. The number of leading basic modes ranged from 3 to 10 (mean ± SD = 5.4 ± 1.39 for REST1 and mean ± SD = 5.5 ± 1.39 for REST2). HCP, Human Connectome Project.

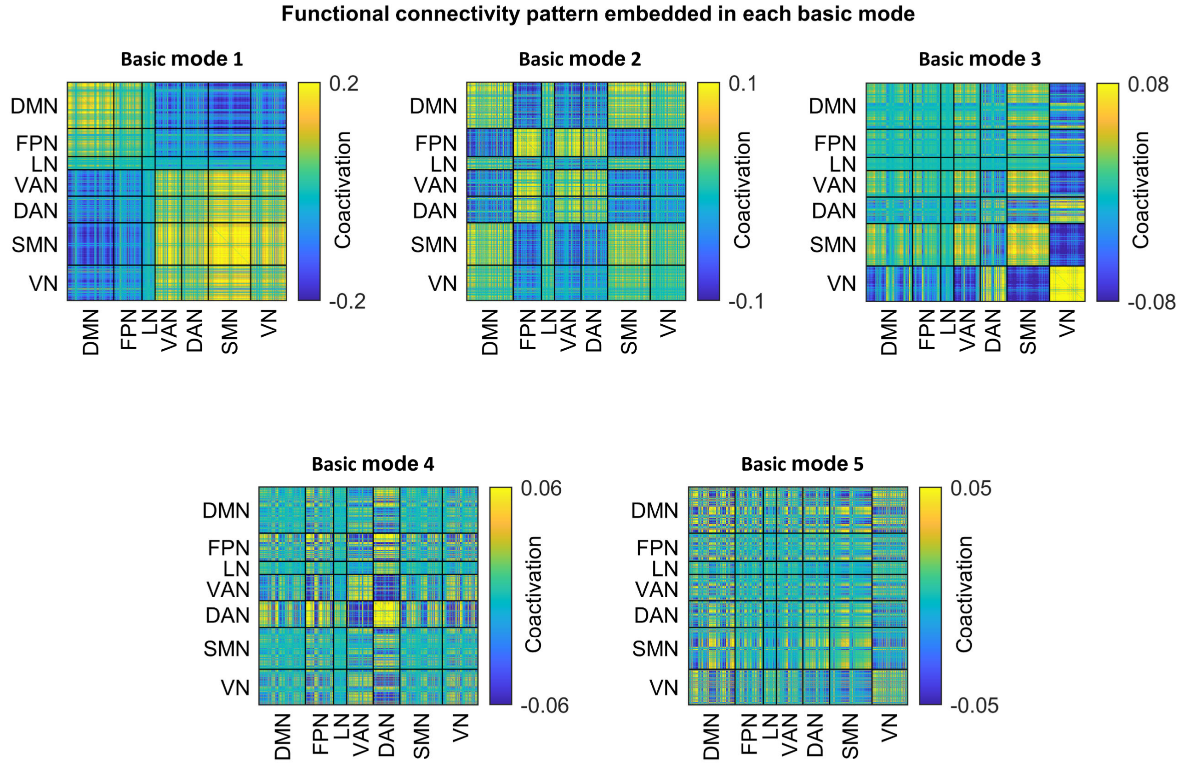


**Figure S3.** Functional connectivity patterns corresponding to five leading basic modes (REST1 of the HCP dataset). For each basic mode, the functional connectivity between two regions was defined as the inter-regional coactivation pattern embedded in the basic mode. The full-scan FC patterns can be regarded as the superposition of these basic FC patterns, suggesting multiplexed relationships are simultaneously present between the same pairs of regions. Nodes are ordered according to their affiliations to prior seven functional systems[1]. DMN, default-mode network; FPN, frontoparietal network; LN, limbic network; VAN, ventral attention network; DAN, dorsal attention network; SMN, somatomotor network; VN, visual network; HCP, Human Connectome Project.

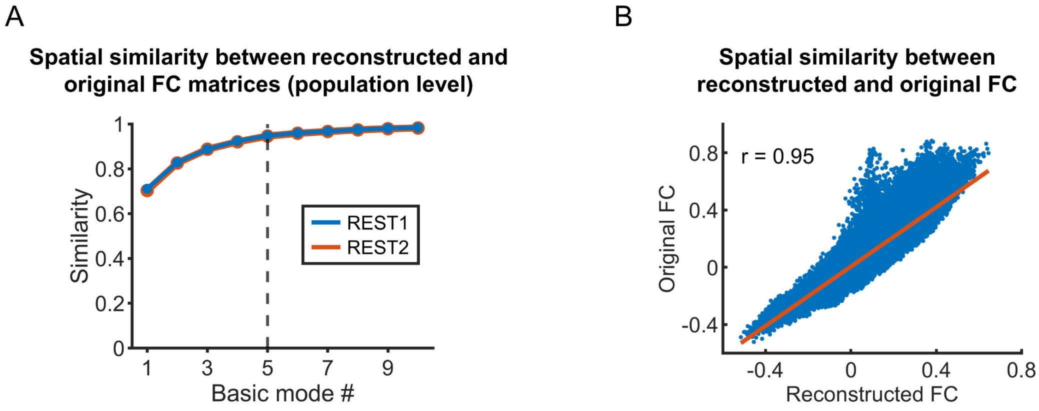


**Figure S4.** Reconstruction of functional connectivity based on the basic modes at the population level. (A) Spatial similarity between the reconstructed and original FC matrices at the population level for both runs. The original FC matrix was estimated based on the concatenated time courses of all participants. The reconstructed FC matrix was generated separately by using different numbers of basic modes. (B) Spatial similarity between the reconstructed and original FC matrices for REST2. The reconstructed FC was generated by using five leading basic modes. The spatial similarity was estimated as Pearson's correlation across the lower triangular elements between two matrices. FC, functional connectivity; HCP, Human Connectome Project.

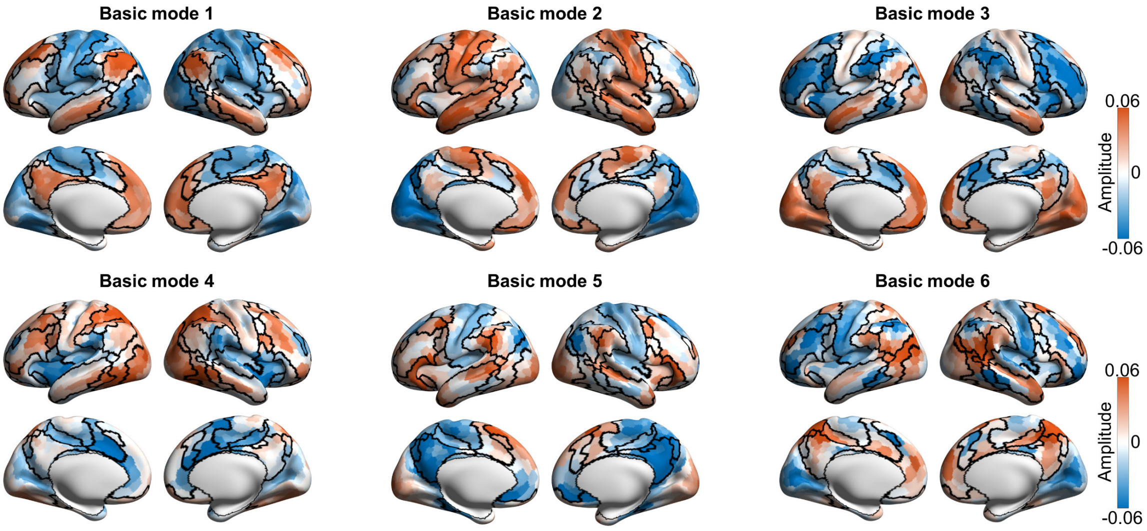


**Figure S5.** Spatial patterns of six leading basic modes (i.e., leading basic modes) at rested wakefulness state in the sleep-deprivation dataset. Black curves denote the boundaries of seven prior functional systems[1].

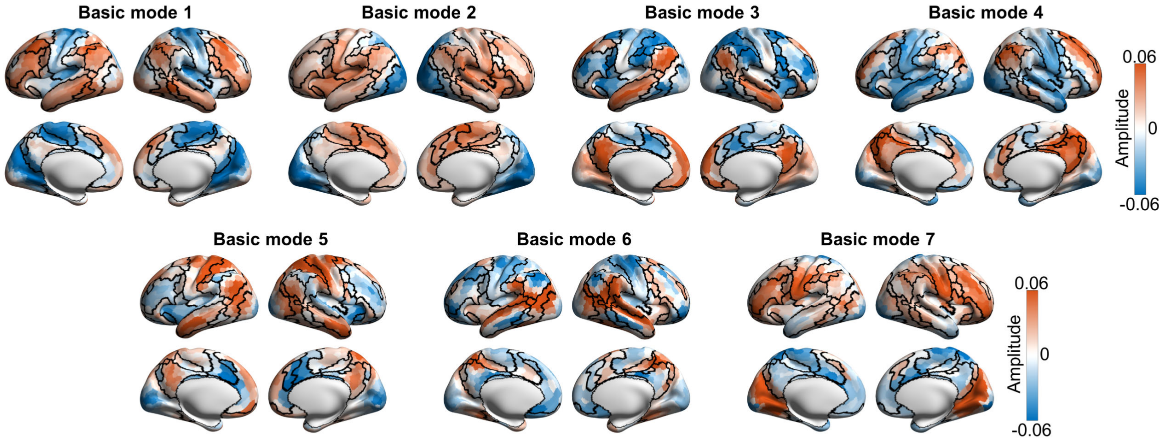


**Figure S6.** Spatial patterns of seven leading basic modes after sleep-deprivation in the sleep-deprivation dataset. Black curves denote the boundaries of seven prior functional systems[1].

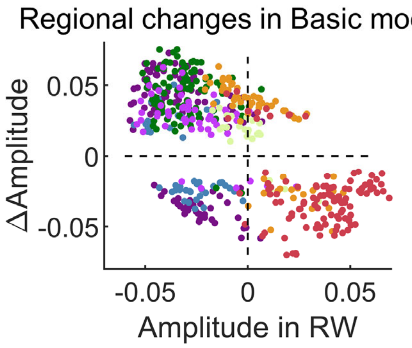


**Figure S7.** Relationship between sleep deprivation-induced changes and the original fluctuation amplitudes at rested wakefulness (for basic mode 1). Only regions with significant changes are displayed. For most regions, the directions of the amplitude changes were opposite to the signs of the original amplitudes, which were mainly located in the default mode network, the dorsal and ventral attention networks, and the frontoparietal network. Colors of the nodes indicate their affiliations of the function system: Red, default-mode network; orange, frontoparietal network; cream, limbic network; violet, ventral attention network; green, dorsal attention network; blue, somatomotor network; purple, visual network. RW, rested wakefulness.

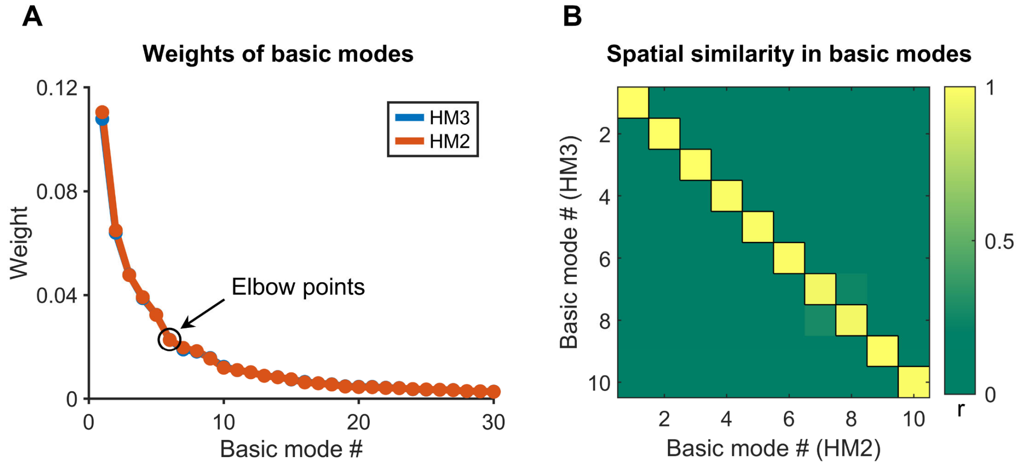


**Figure S8.** Influence of head motion on the identification of leading basic modes (REST1 of the HCP dataset). (A) Weights of basic modes for R-fMRI data screened by two different head motion exclusion criteria (HM3 and HM2). Only the weights of the first thirty basic mode are displayed. Similar decreasing trends were observed in both cases. Elbow points were identified at the sixth basic mode for both runs. (B) Spatial similarity of basic modes between two cases (HM3 and HM2). We found the basic modes were exactly matched between two cases, indicating a weak influence of head motion. With the stricter exclusion criteria, 415 participants (age range: 22-35 years, M/F: 192/223) were retained for further analysis. HM3: the exclusion criterion used in the main result, where participants with head motion greater than 3 mm or 3° in any direction or mean FD greater than 0.5 mm were excluded. HM2: the stricter exclusion criterion, where participants with head motion greater than 2 mm or 2° in any direction or mean FD greater than 0.2 mm were excluded. FD, framewise displacement; HCP, Human Connectome Project.

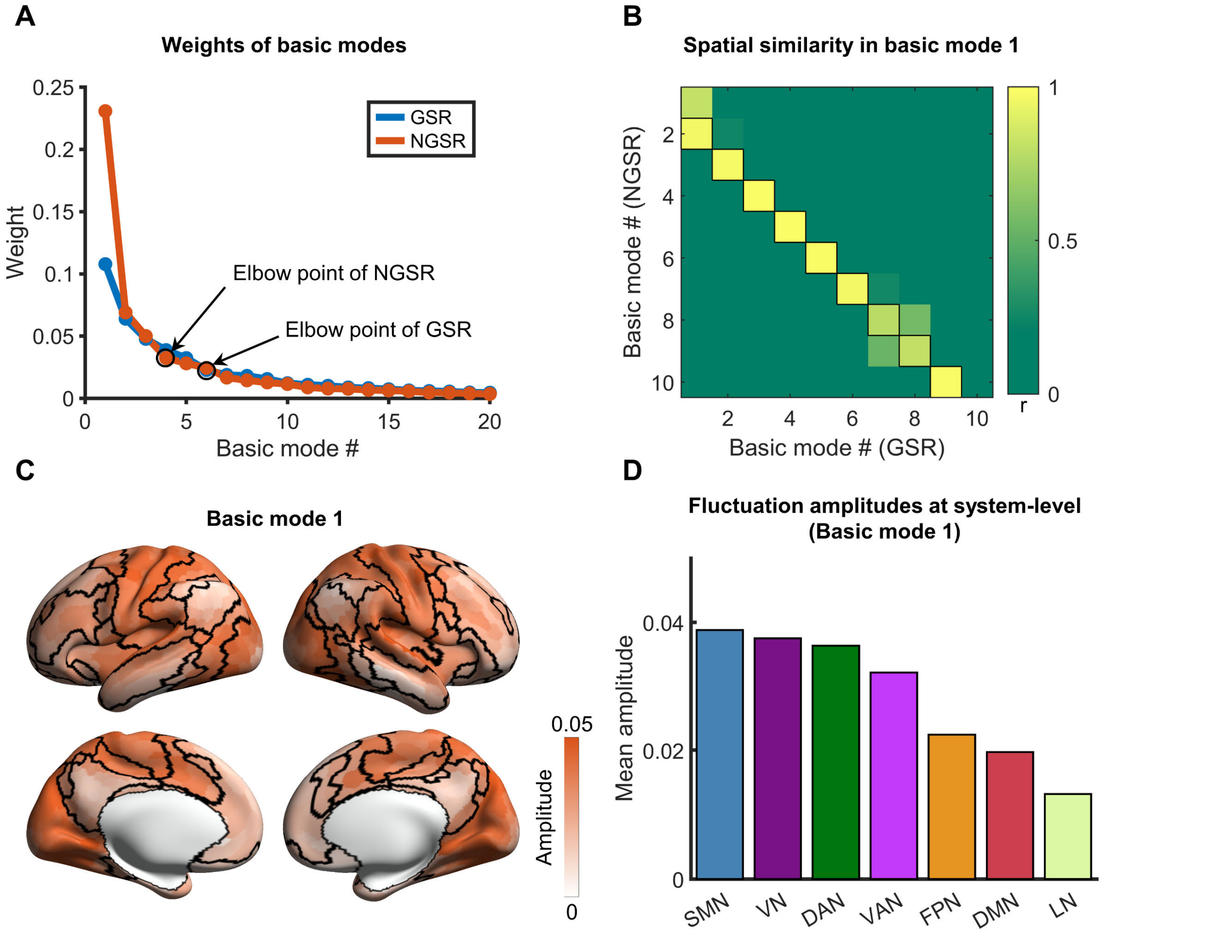


**Figure S9.** Influence of global signal regression on the identification of leading basic modes (REST1 of the HCP dataset). (A) Weights of basic modes with or without global signal regression. Only the weights of the first thirty basic modes are displayed. Note that the weight of the first basic mode obtained from the NGSR condition was much higher than those of the other basic modes. Similar decreasing trends were observed when matching the *i*th basic mode in the GSR condition with the *i*+1th basic mode in the NGSR condition. The elbow points were identified at the fifth basic mode in the NGSR condition. (B) Spatial similarity of basic modes between two cases (GSR and NGSR). We found that the *i*th basic mode in the GSR condition showed the highest spatial similarity with the *i*+1th basic mode in the NGSR condition, suggesting a shift of the basic mode. (C) Spatial pattern of the first basic mode in the NGSR condition. All the brain regions showed the same sign of fluctuation amplitude, suggesting enhanced coactivation across the brain. (D) System-level fluctuation amplitudes for the first basic mode in the NGSR condition. Seven prior functional systems[1] were used here. The ranking of functional systems was similar to that of the first leading basic mode in the GSR condition but in a reversed order. HCP, Human Connectome Project; GSR: with global signal regression; NGSR: without global signal regression; SMN, somatomotor network; VN, visual network; DAN, dorsal attention network; VAN, ventral attention; FPN, frontoparietal network; DMN, default-mode network; LN, limbic network.

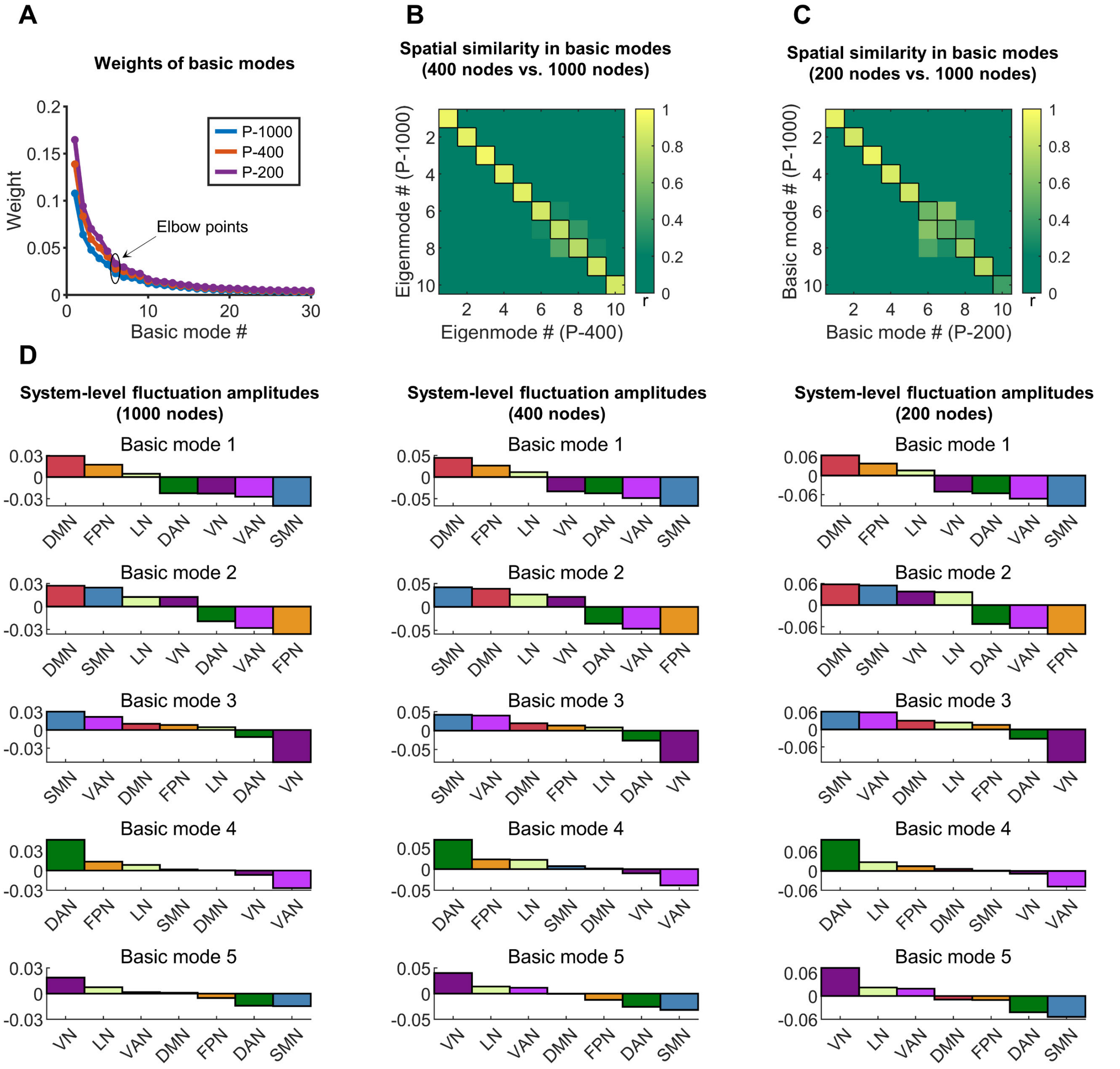


**Figure S10.** Influence of brain parcellations on the identification of leading basic modes (REST1 of the HCP dataset). (A) Weights of basic modes identified with brain parcellations with different spatial resolutions. Three functional parcellations of the cortical cortex were considered, which consisting of 1000 nodes, 400 nodes, and 200 nodes respectively. Only weights of the first thirty basic modes are displayed. Similar decreasing trends were observed for the analyses with different parcellations. Notably, five leading basic modes were identified, regardless of the parcellations. The total weight explained by the five leading basic modes increased with decreasing spatial resolution (29%, 37%, and 44% for the 1000-node, 400-node and 200-node parcellations, respectively). (B) Spatial similarity of the top 10 basic modes between the 1000-node parcellation and the 400-node parcellation. Exact correspondence was observed for all the ten basic modes. (C)

Spatial similarity of the top 10 basic modes between the 1000-node parcellation and 200-node parcellation. Exact correspondence was observed for the five leading basic modes. In (B) and (C), we analyzed the spatial similarity in the basic modes between different parcellations at the voxel level. (D) System-level fluctuation amplitudes for different parcellations. Seven prior functional systems[1] were used here. For each basic mode, similar ranking of the functional systems was observed across three functional parcellations. HCP, Human Connectome Project; P-1000, parcellation with 1000 nodes; P-400, parcellation with 400 nodes; P-200, parcellation with 200 nodes; DMN, default-mode network; FPN, frontoparietal network; LN, limbic network; VAN, ventral attention network; DAN, dorsal attention network; SMN, somatomotor network; VN, visual network.

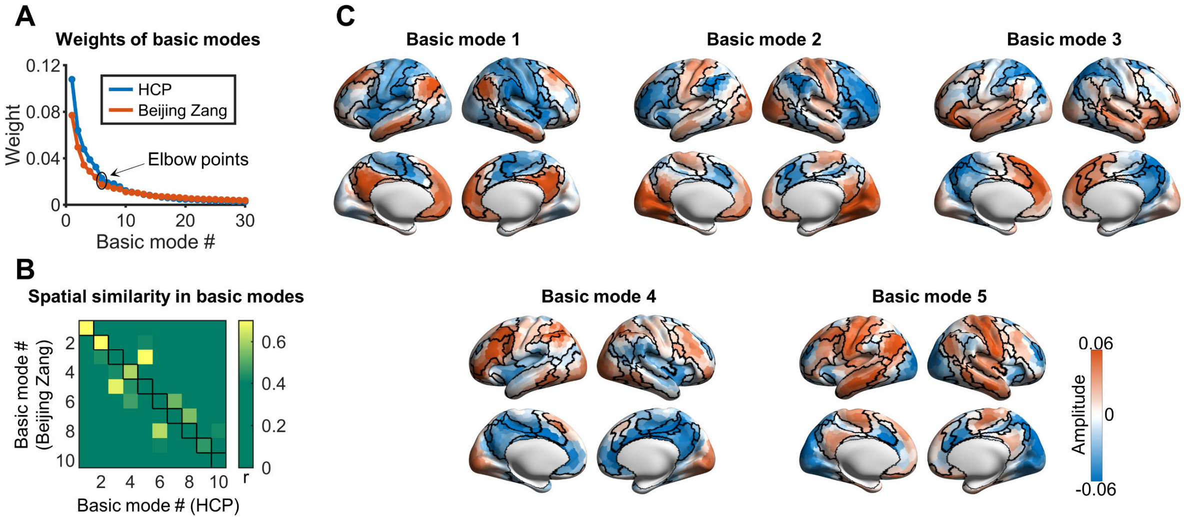


**Figure S11.** Reproducibility of five leading basic modes in an independent dataset (the Beijing Zang dataset). (A) Weights of basic modes for different datasets (i.e., REST1 of the HCP dataset and the Beijing Zang dataset). Only the weights of the first thirty basic modes are displayed. Similar decreasing trends of the weights were observed for both datasets. Similar to the HCP dataset, five leading basic modes were identified for the Beijing Zang dataset. (B) Spatial similarity of the basic modes for both datasets. The first five basic modes were spatially matched, except for the inversion of the 3rd and 5th basic modes. These results suggest that each of the five leading basic modes identified from the main results showed a spatial correspondence in an independent dataset. (C) Spatial patterns of the five leading basic modes for the Beijing Zang dataset. Black curves denote the boundaries of the prior seven functional systems[1].

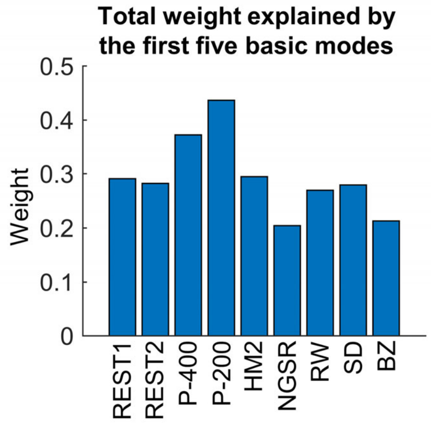


**Figure S12.** Total weight explained by the first five basic modes in different cases. In the NGSR condition, the 2nd to 6th basic modes were considered due to their spatial correspondence to the five leading basic modes in the main results. The first five basic modes accounted for a large proportion of weights (i.e., activity variance) in all cases. Notably, the total weight was parcellation dependent, which increased with decreasing spatial resolution. HCP, Human Connectome Project; P-200, REST1 in the HCP dataset with 200 node parcellation; P-400, REST1 in the HCP dataset with 400 node parcellation; RW, rested wakefulness; SD, after sleep-deprivation; HM2, the stricter exclusion criterion of head motion; NGSR, without global signal regression; BZ, the Beijing Zang dataset.
